# Mutation spectrum of *NOD2* reveals recessive inheritance as a main driver of Early Onset Crohn’s Disease

**DOI:** 10.1101/098574

**Authors:** Julie E. Horowitz, Neil Warner, Jeffrey Staples, Eileen Crowley, Ryan Murchie, Cristopher Van Hout, Alejandra Klauer King, Karoline Fiedler, Jeffrey G. Reid, John D. Overton, Alan R. Shuldiner, Aris Baras, Geisinger-Regeneron DiscovEHR Collaboration, Frederick E. Dewey, Anne Griffiths, Omri Gottesman, Aleixo M. Muise, Claudia Gonzaga-Jauregui

## Abstract

Inflammatory bowel disease (IBD), clinically defined as Crohn’s disease (CD), ulcerative colitis (UC), or IBD-unclassified, results in chronic inflammation of the gastrointestinal tract in genetically susceptible hosts. Pediatric onset IBD represents ≥25% of all IBD diagnoses and often presents with intestinal stricturing, perianal disease, and failed response to conventional treatments. *NOD2* was the first and is the most replicated locus associated with adult IBD, to date. To determine the role of *NOD2* and other genes in pediatric IBD, we performed whole-exome sequencing on a cohort of 1,183 patients with pediatric onset IBD (ages 0-18.5 years). We identified 92 probands who were homozygous or compound heterozygous for rare and low frequency *NOD2* variants accounting for approximately 8% of our cohort, suggesting a Mendelian recessive inheritance pattern of disease. Additionally, we investigated the contribution of recessive inheritance of *NOD2* alleles in adult IBD patients from the Regeneron Genetics Center (RGC)-Geisinger Health System DiscovEHR study, which links whole exome sequences to longitudinal electronic health records (EHRs) from 51,289 participants. We found that ~7% of cases in this adult IBD cohort, including ~10% of CD cases, can be attributed to recessive inheritance of *NOD2* variants, confirming the observations from our pediatric IBD cohort. Exploration of EHR data showed that 14% of these adult IBD patients obtained their initial IBD diagnosis before 18 years of age, consistent with early onset disease. Collectively, our findings show that recessive inheritance of rare and low frequency deleterious *NOD2* variants account for 7-10% of CD cases and implicate *NOD2* as a Mendelian disease gene for early onset Crohn’s Disease.

**Author Summary:** Pediatric onset inflammatory bowel disease (IBD) represents ≥25% of IBD diagnoses; yet the genetic architecture of early onset IBD remains largely uncharacterized. To investigate this, we performed whole-exome sequencing and rare variant analysis on a cohort of 1,183 pediatric onset IBD patients. We found that 8% of patients in our cohort were homozygous or compound heterozygous for rare or low frequency deleterious variants in the nucleotide binding and oligomerization domain containing 2 *(NOD2)* gene. Further investigation of whole-exome sequencing of a large clinical cohort of adult IBD patients uncovered recessive inheritance of rare and low frequency *NOD2* variants in 7% of cases and that the relative risk for *NOD2* variant homozygosity has likely been underestimated. While it has been reported that having >1 *NOD2* risk alleles is associated with increased susceptibility to Crohn’s Disease (CD), our data formally demonstrate what has long been suspected: recessive inheritance of *NOD2* alleles is a mechanistic driver of early onset IBD, specifically CD, likely due to loss of NOD2 protein function. Our data suggest that a subset of IBD-CD patients with early disease onset is characterized by recessive inheritance of *NOD2* alleles, which has important implications for the screening, diagnosis, and treatment of IBD.

## Introduction

Inflammatory bowel disease (IBD) is a chronic inflammatory condition of the gastrointestinal (GI) tract that arises as part of an inappropriate response to commensal or pathogenic microbiota in a genetically susceptible individual (1–4). IBD encompasses Crohn’s Disease (CD); ulcerative colitis (UC); and IBD unclassified (IBDU). The etiology of IBD is complex and has been attributed to defects in a number of cellular pathways including pathogen sensing, autophagy, maintenance of immune homeostasis, and intestinal barrier function, among other processes (3–18).

Great effort has been invested into defining the genetic factors that confer IBD susceptibility. To date, >200 unique loci have been associated with IBD through genome-wide association studies (GWAS), primarily in adult populations (19, 20). Nearly all the identified susceptibility loci exhibit low effect sizes (ORs ~1.0–1.5) individually (19), and collectively account for less than 20% of the heritable risk for IBD (19, 21). These observations support a complex disease model in which common variants of modest effect sizes interact with environmental factors including diet, smoking, and the intestinal microbiome (22, 23) to give rise to IBD susceptibility (4).

The earliest and most replicated genetic associations with IBD (24–26) correspond to a locus on chromosome 16 that encompasses the nucleotide-binding and oligomerization domain-containing 2 *(NOD2)* gene, with an average allelic odds ratio across multiple studies of *3.1* (19, 20). *NOD2* encodes an intracellular microbial sensor that recognizes muramyl dipeptide (MDP) motifs found on bacterial peptidoglycans (27, 28). Upon activation, NOD2 protein signals through the NF-κB family of proteins (29) to modulate transcription of genes encoding pro-inflammatory cytokines IL-8, TNF-α, and IL-1 β (30–32), among others. Variation in *NOD2* accounts for approximately 20% of the genetic risk among CD cases, with three variants - p.R702W (ExAC MAF= 0.0227 across all populations), p.G908R (ExAC MAF= 0.0099), and p.L1007fs (ExAC MAF= 0.0131) - accounting for over 80% of the disease-causing mutations in *NOD2* associated with adult CD (33), albeit not with UC; and particularly ileal versus colonic CD (34). These three “common” risk variants, typically observed in a heterozygous state, are predicted loss-of-function alleles that impair NF-κB activation in response to MDP ligands, *in vitro (28, 35–37).*

With the assumption that genetic risk has a disproportionate effect over environmental risk in early onset disease, recent studies have focused on pediatric IBD cases (diagnosed <18y) (38). Pediatric IBD patients comprise 20-25% of all IBD cases and are typically more clinically severe than adult-onset patients, often exhibiting disease of the upper GI tract, small bowel inflammation, and perianal disease as well as failure to thrive and poor clinical response (4, 39). Results from GWAS conducted in this group of severely affected patients indicate that associated loci in early onset IBD significantly overlap with adult IBD loci, including both the *NOD2* locus and an additional 28 CD-specific loci previously implicated in adult-onset IBD (40–42). As the mechanism for these “common” IBD susceptibility loci in the pathogenesis of early onset IBD remains unclear (43), we performed whole-exome sequencing and rare variant analysis on a cohort of 1,183 pediatric onset IBD patients to elucidate the role of rare protein coding variation in IBD-associated genes, specifically *NOD2,* in this disease.

## Results and Discussion

We performed trio-based analysis of 492 complete trios using a proband-based analytical pipeline to identify all recessive (compound heterozygous and homozygous) and *de novo* variants of interest in the affected probands. In our initial analyses, we identified 10 families with recessive (compound heterozygous or homozygous), rare variants (2%≤MAF) in *NOD2,* all with a diagnosis of CD. We observed that some of the rare variants in these probands were inherited in *trans* from previously-reported CD risk alleles, mainly the p.G908R missense variant. We identified two individuals who are compound heterozygous for the p.G908R risk allele in *trans* with a less common *NOD2* CD risk variant (p.N852S) in one case and a novel truncating indel (p.S506Vfs*73) in the second case (Table S1, Fam008 and Fam009). The observation of a CD-associated *NOD2* risk allele in *trans* from other rare or novel alleles led us to survey the rest of the probands, including singletons and those part of incomplete trios, for recessive inheritance, either in a homozygous or compound heterozygous manner, of *NOD2* variants, but expanding our allelic range to low-frequency variants (2%≤MAF≤5%). Through this approach we identified 108 probands with putative recessive *NOD2* variants. Visual inspection of sequence reads and orthogonal confirmation through Sanger sequencing excluded 13 probands with variants inherited in *cis* from an unaffected parent or heterozygous variants that were initially called as homozygous due to low coverage of the region and skewed allelic balance. Of note, we identified 5 probands carrying p.L1007fs and p.M863V risk variants, 4 of which were confirmed to occur in *cis* and were inherited from an unaffected parent. The remaining case with p.L1007fs and p.M863V was a singleton and thus phase could not be determined. These two variants segregate in *cis* within the same haplotype, as confirmed by segregation within the trios and as previously observed (44). Therefore, we excluded these 4 probands from our final count of recessively-inherited *NOD2* variants. Similarly, we identified 3 probands from 3 complete trios segregating the p.S431L and p.V793M reported risk variants in *cis* inherited from an unaffected carrier parent; these probands were also excluded. Three additional probands were excluded on the basis of a re-evaluation of the phenotype that excluded a clinical diagnosis of IBD.

Thus, we identified 92 probands with confirmed recessive *NOD2* variants within our pediatric onset IBD cohort. These include: 25 probands carrying homozygous variants, 41 probands with confirmed compound heterozygous variants, and an additional 26 singleton probands with putative compound heterozygous variants where phasing could not be performed (Table S1, Figure S1). The majority of the compound heterozygous individuals (65/67) carry a known *NOD2* CD-risk allele in addition to either another known *NOD2* CD-risk allele or a novel *NOD2* variant, including some truncating loss-of-function variants supporting loss or impaired function of NOD2 in the pathophysiology of CD (6). In total, 92 of 1,183 (7.8%) of the probands in our pediatric onset IBD cohort conformed to a recessive, Mendelian inheritance mode for *NOD2* rare and low frequency (MAF≤5%) deleterious variants (Table 1, Figure 1, Table S1, and Table S4).

**Figure 1.**
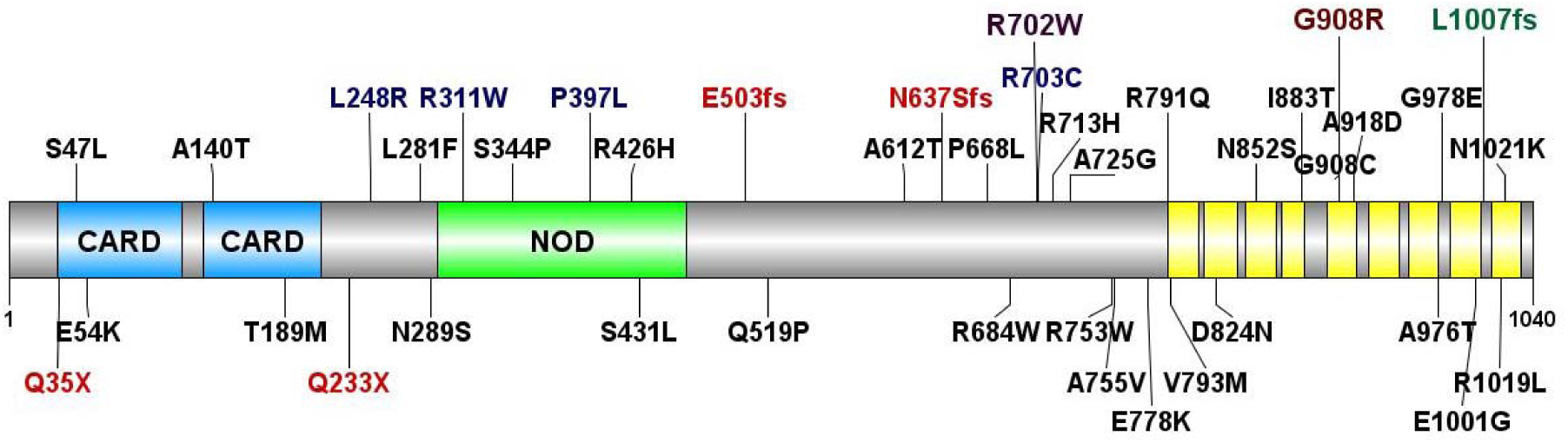
Mutation spectrum of *NOD2* in inflammatory bowel disease (IBD) patients. *NOD2* variation identified in patients with pediatric early onset IBD (upper) and adult IBD cohort from the RGC-GHS DiscovEHR collaboration (lower). Variants in blue were observed in both cohorts; variants in red are predicted loss-of-function that result in nonsense mediated decay. The three “common” low-frequency Crohn’s Disease risk variants are highlighted: R702W (purple), G908R (brown), and L1007fs (green). Also depicted are the NOD2 protein structural domains: two caspase activation and recruitment domains (CARD), a nucleotide binding and oligomerization (NOD) domain, and leucine rich repeat domains in yellow.

**Table 1.** Mutation spectrum of recessive *NOD2* variants in an EO-IBD cohort.

| NOD2 Variant (p.) | # EO-IBD Probands | Mean Age (Range) | % CD Dx | Tissue Involvement |  |  |
| --- | --- | --- | --- | --- | --- | --- |
|  |  |  |  | <u>Colon</u> | <u>Ileum</u> | <u>Perianal</u> |
| <u>Compound Heterozygous</u> |  |  |  |  |  |  |
| Rare/Rare | 4T; 1S | 12.5 (9.0-14.6) | 88.9% | 80.0% | 60.0% | 40.0% |
| Common/Rare | 1Q; 9T; 6D; 14S | 12.6 (5.5-16.5) | 93.1% | 57.1% | 82.1% | 25.0% |
| Common/Common | 17T; 5D; 10S | 11.8 (2.1-18.5) | 90.6% | 56.3% | 90.6% | 25.0% |
| <u>Homozygous</u> |  |  |  |  |  |  |
| Rare | 1D; 2S | 9.6 (4.2-15.1) | 66.7% | 33.3% | 33.3% | 33.3% |
| p.R702W | 1Q; 2T; 3D; 1S | 11.7 (5.8-13.8) | 100% | 85.7% | 57.1% | 42.9% |
| p.G908R | 2T; 1D; 2S | 10.6 (7.7-13.7) | 80.0% | 40.0% | 40.0% | 0.0% |
| p.L1007fs | 4T; 2D; 4S | 12.1 (5.9-13.4) | 100% | 60.0% | 90.0% | 20.0% |

Common *NOD2* variants refer to the three main low-frequency Crohn's Disease risk variants p.R702W, p.G908R, and p.L1007fs; Rare *NOD2* variants refer to other low-frequency variants ( $MAF \leq 5\%$ ). Abbreviations: Q, quartet; T, trio; D, duo; S, singleton; Dx, diagnosis

The 92 pediatric patients homozygous for *NOD2* mutations were predominantly male (71%) with a median age at diagnosis of 12.5 years (Table S1). At diagnosis, 83% displayed diagnostic features of Crohn’s disease. 23% of the cohort displayed a constellation of extra-intestinal manifestations, mainly large joint arthritis, chronic recurrent multifocal osteomyelitis, recurrent fevers, erythema nodosum, and pyoderma gangrenosum. Only 6% of the cohort showed significant perianal disease (namely fistulae and abscesses, skin tags and fissures were not considered as perianal disease) (Table S1). Per the Montreal classification of IBD (45), 44% of the overall cohort of patients presented with ileal disease at diagnosis (L1). 25% presented with ileocolonic disease (L3) and 10% displayed features of colonic inflammation only (L2). Isolated upper disease was only present in 2% of the cohort (L4). We observed a progression of ileal disease in 21% (18.5% stricturing; 2.5% penetrating) with 21% requiring a resection. On review of the Crohn’s disease patients only, 50% displayed L1 disease (terminal ileal +/- limited cecal disease), 32.9% L3 (ileocolonic) disease; 86.8% B1 (non-stricturing, non-penetrating) disease, 10.5% B2 (stricturing) disease (Table S1).

Given the substantial contribution of recessive *NOD2* variants to CD in our pediatric onset IBD cohort and the known contribution of *NOD2* to adult CD, we next investigated the contribution of *NOD2* recessivity in a large clinical population. For this, we examined a cohort of adult IBD patients from the RGC - Geisinger Health System (GHS) DiscovEHR study (46) . A key feature of the DiscovEHR study is the ability to link genomic sequence data to de-identified electronic health records (EHRs). Within this cohort, we identified 984 patients (of 51,289 total sequenced DiscovEHR patient-participants) with a diagnosis of IBD, defined as having a problem list entry or an encounter diagnosis entered for two separate clinical encounters on separate calendar days for the ICD-9 codes 555* (Regional enteritis) or 556* (Ulcerative enterocolitis). For our analysis, we surveyed all instances of homozygous *NOD2* rare and low frequency variants (MAF≤5%); the same parameters applied to our pediatric IBD probands. Among patients with an IBD diagnosis, we identified 18 individuals who are either homozygous for the p.R702W risk allele (N=10) or homozygous for the p.L1007fs allele (N=8) (Table 2, Figure S2). We did not identify any p.G908R homozygous individuals with an IBD diagnosis in this cohort. Next, we looked for instances of putative compound heterozygosity among these adult IBD DiscovEHR patients. To investigate this, we searched for occurrences of two or more of the three most prevalent *NOD2* risk alleles (p.R702W, p.G908R, or p.L1007fs) in these individuals. We identified putative compound heterozygosity for the three main CD risk alleles, p.R702W/p.G908R (N=6), p.G908R/p.L1007fs (N=5), and p.R702W/p.L1007fs (N=11) (Table 2). We also observed instances of putative compound heterozygosity for each of the three main CD risk alleles along with either a rarer CD risk allele or a novel allele or two rare alleles in *trans* (N=24), parallel to the findings in our pediatric IBD cohort. Using familial relationships and pedigree reconstruction (47), we were able to confirm appropriate segregation for 32 of the 64 DiscovEHR recessive *NOD2* variant carriers with IBD, including *trans* inheritance in 13 putative compound heterozygotes (Figure S2). The other 32 were singleton cases where phase could not be confirmed. Overall, we identified 64 homozygous or putative compound heterozygous *NOD2* variant carriers in the DiscovEHR IBD cohort, accounting for 6.5% of patients with an IBD diagnosis in this clinical population (Figure 1, Table S4).

**Table 2.** Mutation spectrum of recessive *NOD2* variants in the RGC-GHS DiscovEHR adult IBD cohort.

| NOD2 Variants | # IBD patients | Mean Age (Range) | # CD Dx | % CD Dx | # Male | % Male | # Perianal | % Perianal |
| --- | --- | --- | --- | --- | --- | --- | --- | --- |
| <u>Homozygous</u> |  |  |  |  |  |  |  |  |
| p.R702W | 10 | 47.3 (16.0-76.3) | 10 | 100% | 5 | 50.0% | 1 | 10.0% |
| p.G908R | 0 | - | - | - | - | - | - | - |
| p.L1007fs | 8 | 40.25 (11.0-75.0) | 8 | 100% | 4 | 50.0% | 3 | 37.5% |
| <u>Compound Heterozygous</u> |  |  |  |  |  |  |  |  |
| <u>Common/Common</u> |  |  |  |  |  |  |  |  |
| p.R702W/p.G908R | 6 | 48.5 (11.4-54.2) | 3 | 50.0% | 5 | 83.3% | 1 | 16.7% |
| p.R702W/p.L1007fs | 11 | 38.5 (21.5-69.2) | 8 | 72.7% | 3 | 27.3% | 2 | 18.2% |
| p.G908R/p.L1007fs | 5 | 35.2 (20.0-52.4) | 4 | 80.0% | 2 | 40.0% | 0 | 0.0% |
| <u>Common/Rare</u> | 16 | 40.3 (10.8-66.1) | 8 | 50.0% | 8 | 50.0% | 6 | 37.5% |
| <u>Rare/Rare</u> | 8 | 60.9 (30.8-78.7) | 4 | 50.0% | 3 | 37.5% | 1 | 12.5% |

Common *NOD2* variants refer to the three main low-frequency Crohn's Disease risk variants p.R702W, p.G908R, and p.L1007fs; Rare *NOD2* variants refer to other low-frequency variants ( $MAF \leq 5\%$ ). Abbreviations: Dx, diagnosis.

We were also able to evaluate longitudinal de-identified medical records for all patients within the DiscovEHR IBD cohort. According to their EHR data, 21 patients received diagnoses of both UC and CD. To clarify these diagnoses, we performed an evaluation of EHR information (which includes demographics, encounter and problem list diagnosis codes, procedure codes, and medications) for all 64 homozygous or compound heterozygous *NOD2* patients with an IBD diagnosis. Through this review, 6 homozygotes exhibited a conflicting diagnosis of CD, of which 5 were resolved as CD and 1 could not be defined; 16 compound heterozygotes exhibited a conflicting diagnosis of CD of which 6 were resolved as CD and 10 were resolved as UC (Table S3). In total, we found that 17/18 (94.4%) of homozygous *NOD2* individuals and 33/46 (71.7%) compound heterozygous had a diagnosis of CD and that 9.9% of all CD cases in this cohort could be attributed to homozygous or compound heterozygous variants in *NOD2.* We next investigated age of disease onset using the first recorded date of an IBD diagnosis in the EHR. We identified 6 carriers of recessive *NOD2* variants (9.4% of our recessive *NOD2* patients with IBD) who were diagnosed with IBD prior to 18 years of age. We also identified an additional 11 carriers of recessive *NOD2* variants diagnosed with IBD prior to age 30 years, which is at or below the average age of IBD diagnosis (48) and is consistent with earlier disease onset (Table S3). Of note, our RGC-GHS DiscovEHR data extends to a median of 14 years (and maximum of 20 years) of electronically recorded medical information, concurrent with the adoption of the EHR by the Geisinger Health System. Since 72.4% of our cohort is currently over the age of 50 years, we cannot determine whether the age of onset for IBD occurred prior to the first electronically recorded date of an IBD diagnosis for many recessive *NOD2* patients; thus it is possible that other individuals with homozygous or compound heterozygous variants in *NOD2* might have had pediatric-onset disease that was not captured in the EHR.

Finally, given the recessive inheritance of *NOD2* variants observed in both our pediatric onset and adult IBD cohorts, we estimated the disease risk for the three main known CD risk alleles (p.R702W, p.G908R, and p.L1007fs) in our adult IBD case cohort and their effect sizes using additive, genotypic, and recessive genetic models. Under an additive model, we observed similar effect sizes for each of the 3 variants [OR=1.43 (1.20-1.71 95%CI, P-value 4.63×10^−5^) for p.R702W; OR=1.56 (1.18–2.06 95%CI, P-value 1.54×10^−3^) for p.G908R; and OR=1.84 (1.50–2.26 95%CI, P-value 1.69×10^−9^) for p.L1007fs], consistent with previously reported low to moderate effect sizes for each allele by GWAS (19) and the most recent data available in the IBD Exomes Portal (49) (Table 3, Figure 2). However, for the two risk alleles with homozygous cases, in the genotypic model – which estimates distinct effect sizes for heterozygous and homozygous carriers – we observe substantially larger effects in homozygotes versus heterozygotes for the p.R702W variant (Het OR= 1.30 [1.06–1.58 95%CI], P-value 8.77×10^−3^, versus Hom OR= 4.02 [2.17-7.45 95% CI], P-value 6.86X10^−6^) and the p.L1007fs variant (Het OR= 1.63 [1.30–2.04 95%CI], P-value 1.67×10^−5^, versus Hom OR= 10.15 [4.75-21.69 95% CI], P-value 1.38×10^−12^). We also calculated the effect sizes using a recessive model for these two variants and the 22 compound heterozygotes carrying any combination of the 3 CD risk alleles. We found that recessive effect sizes for the p.R702W and p.L1007fs variants were similar to those observed under the homozygous genotypic model (OR=3.91 [2.11–7.24 95% CI], P-value 2.86×10^−6^, and OR=9.81 [4.59–20.94 95% CI], P-value 3.80X10^−13^, respectively) (Table 3, Figure 2). Further, under the recessive model we observed that the effect size for the compound heterozygotes was also significant (OR=4.35 [2.80–6.75 95% CI], P-value= 8.14×10^−13^), consistent with our previous observations (Table 3, Figure 2). Finally, we calculated the combined contribution of the 3 CD risk alleles under the different genetic models: additive (OR=1.64 [1.45-1.86 95%CI], P-value 4.58×10^−15^), genotypic (Het OR= 1.49 [1.28–1.73 95%CI], P-value 2.75×10^−7^, versus Hom OR= 5.24 [3.77-7.27 95% CI], P-value 4.31X10^−22^), and recessive (OR=4.81 [3.47–6.67 95% CI], P-value= 1.63×10^−25^) (Table 3, Figure 2). Collectively, these analyses show substantially larger effects for *NOD2* homozygotes and compound heterozygotes than heterozygotes and indicate that the genetic contribution of *NOD2* alleles, in a subset of IBD patients, is consistent with a recessive disease model.

**Figure 2.**
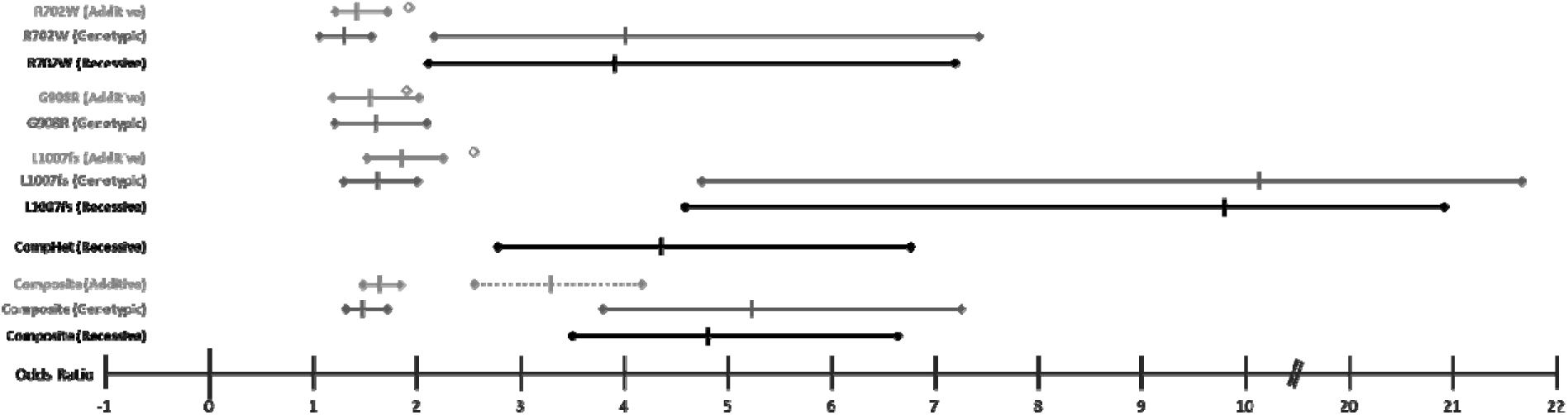
Graphical representation of Odds Ratio (OR) point estimates and 95% confidence intervals (CI) for the three main CD risk alleles (p.R702W, p.G908R, p.L1007fs) under additive, genotypic, and recessive genetic models. (corresponding to values in Table 3). The dotted line in the Composite panel depicts the calculated CI with corresponding calculated OR for 2 alleles under an additive genetic model; of note the point estimate (2xOR) is outside of the 95% CI for the Composite genotypic homozygous and recessive models. Diamonds correspond to estimated OR values for these same variants in the IBD Exomes Browser(49); no confidence intervals are provided.

**Table 3.**
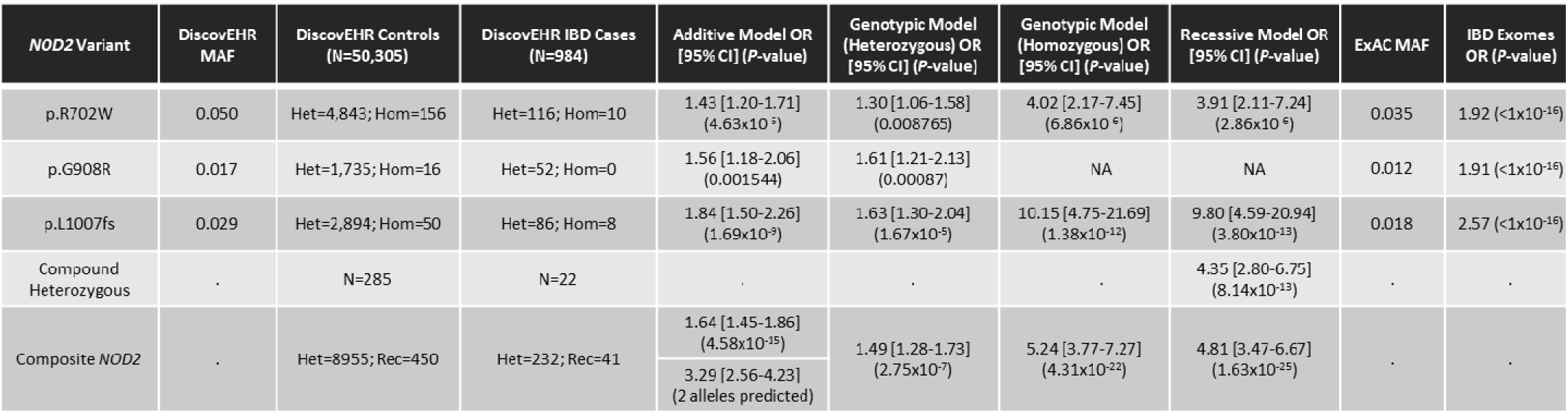
OR calculations for 3 NOD2 CD risk alleles (p.R702W, p.G908R, p.L1007fs), composite, and compound heterozygous combinations in the DiscovEHR cohort. There were no homozygotes for the p.G908R variant affected with IBD in our cohort; therefore, no genotypic homozygous and recessive ORs could be calculated. The ‘Composite *NOD2’* calculations account for all alleles and genotypes for the 3 CD risk variants in the different genetic models.

These observations are in line with previous analyses and meta-analyses of CD cohorts where individuals carrying any one of the main three CD associated risk alleles (p.R702W, p.G908R, or p.L1007fs) have 2-4 fold increased risk for developing CD (36), whereas carriers of two or more of the same *NOD2* variants have a 15-40 fold increased risk for developing CD (33,50,51), exhibiting disease of the terminal ileum (34), and earlier diagnosis (by an average of 3 years) (33). Our observations support these studies but highlight a subset of IBD cases molecularly defined by recessive inheritance of *NOD2* alleles that exhibit markedly increased risk for CD with significantly earlier age of onset (mean age of onset among recessive *NOD2* carriers in the DiscovEHR IBD cohort: 43.4y; mean age of onset in the DiscovEHR IBD cohort: 51.5y; P-value: 4.0X10^−4^ by unpaired t test).

Further, while we observe a low effect size for single allele carriers, based on our allelic effect size calculations for each of the 3 main CD risk alleles in our DiscovEHR cohort (Table 3, Figure 2), we hypothesize that homozygous and compound heterozygous *NOD2* individuals included in large IBD GWAS cohorts have likely contributed to a large proportion of the relative risk calculations for IBD, specifically for CD, under additive models, and that homozygous effect sizes have been largely underappreciated or underreported. It is possible that stratification or conditional statistical analysis of these large and heterogeneous cohorts based on *NOD2* genotypes may increase power to detect other loci that contribute to IBD.

While our observations strongly support recessive inheritance of *NOD2* variants as a driver of early onset Crohn’s disease, we observed incomplete penetrance, as evidenced by homozygous or compound heterozygous *NOD2* variant carriers that do not have a clinical presentation of IBD (52–54). Penetrance and expressivity are two major genetic concepts that play into the onset of the phenotype and the clinical presentation of monogenic diseases (55). In the case of IBD, penetrance is known to be incomplete and clinical presentation is extremely variable. Further, the contribution of additional environmental triggers that may enhance disease onset and/or severity in an already genetically-compromised individual should not be underestimated, especially considering that the loss of epithelial barrier function occurring during IBD allows for host exposure to up to 10^14^ gut microbiota (56, 57). Even in cases of monogenic IBD, such as IL-10 receptor deficiency (58–60), intestinal flora are required for disease presentation in murine disease models (61–63). Furthermore, variation in genes involved in NOD2-dependent signaling pathways, including *XIAP* (64–66) and *TRIM22* (67), result in Mendelian forms of IBD. For *XIAP,* and most likely *TRIM22,* viral triggers are required for disease onset and progression, and *XIAP* mutations have variable penetrance, with only a small percentage of XIAP-deficiency patients developing CD (age of onset between 3 months and 40 years (52)). As NOD2-deficient hosts are more susceptible to the pathogenic effects of a changing intestinal microenvironment (68), the contribution of either discrete or continuous gene-environment exposures may further explain heterogeneity in onset and presentation of disease for genetically-sensitized recessive *NOD2* carriers.

Given the wide variability in clinical presentation of IBD (54), we cannot exclude the possibility that recessive *NOD2* carriers exhibit subclinical phenotypes not formally diagnosed as IBD or that they may eventually develop IBD. It is additionally possible that recessive *NOD2* carriers in the DiscovEHR cohort have a diagnosis of IBD that has not been captured in the EHR. Future investigation into the medical histories of recessive *NOD2* carriers may shed light on this variable expressivity or incomplete capture of medical information. We also cannot exclude the possibility that recessive *NOD2* carriers possess additional genes or alleles that either contribute to disease onset and severity or, alternatively, provide protection or reduced expressivity of the phenotype. Identification of these genetic modifiers warrants future investigation both to unveil additional IBD-risk associated loci for early onset UC and CD cases and to identify protective genes and alleles that can be used to derive therapeutic avenues for IBD treatment and management.

In summary, in a cohort of 1,183 pediatric and early onset IBD patients, we report recessive inheritance of rare and low frequency variants in *NOD2* accounting for about 8% of probands. We assessed the contribution of *NOD2* recessive inheritance in a broader, heterogeneous cohort of adult IBD patients, similar to those recruited for GWAS, and found that recessive inheritance of variants in *NOD2* account for 6.5% of these IBD patients, including 9.9% of CD cases. Thus, recessive inheritance of rare and low frequency *NOD2* variants explain a substantial proportion of CD cases in a pediatric cohort and a large clinical population, with significantly earlier age of disease onset. Consistently, both pediatric and adult CD exhibit a broad spectrum of clinical presentation, suggesting a shared etiology across age groups, at least in the subgroup defined by recessive NOD2-driven CD. Our findings indicate that deleterious *NOD2* variants should be considered as strong predictors of IBD-CD onset and implicate *NOD2* as a Mendelian disease gene for early onset IBD, specifically for a molecularly defined subset of Crohn’s disease patients.

## Materials and Methods

We performed whole exome sequencing on a cohort of 1,183 probands with pediatric onset IBD (ages 0-18.5 years), including their affected and unaffected parents and siblings, where available (total samples = 2,704). All individuals were consented for genetic studies under an IRB-approved protocol by the Toronto Hospital for Sick Children. Sample preparation, whole exome sequencing, and sequence data production were performed at the Regeneron Genetics Center (RGC) as previously described (46). In brief, 1ug of high-quality genomic DNA for exome capture using the NimbleGen VCRome 2.1 design. Captured libraries were sequenced on the Illumina HiSeq 2500 platform with v4 chemistry using paired-end 75 bp reads. Exome sequencing was performed such that >85% of the bases were covered at 20x or greater. Raw sequence reads were mapped and aligned to the GRCh37/hg19 human genome reference assembly, and called variants were annotated and analyzed using an RGC developed pipeline. Briefly, variants were filtered based on their observed minor allele frequencies at a <2% cutoff using the internal RGC database and other population control databases (ExAC (69) and NHLBI’s ESP (70)) to filter out common polymorphisms and high frequency, likely benign variants in consideration of disease prevalence.

## Conflict of Interest Disclosure

J.E.H. is a postdoctoral fellow at the Regeneron Genetics Center, Regeneron Pharmaceuticals, Inc. J.S., C.V.H., A.K.K., J.G.R., J.D.O., A.R.S., A.B., F.E.D, O.G., and C.G.J are full-time employees of the Regeneron Genetics Center, Regeneron Pharmaceuticals, Inc. and receive stock options as part of compensation. N.W., R.M., K.F., A.G., and A.M.M have no conflicts to disclose.

## Acknowledgements

We are thankful to the patients and families who participated in this study. A.M.M. is funded by a CIHR – Operating Grant (MOP119457) and the Leona M. and Harry B. Helmsley Charitable Trust to study VEOIBD.

